# Recruitment and disruption of value encoding during alcohol seeking

**DOI:** 10.1101/513911

**Authors:** David Ottenheimer, Karen Wang, Alexandria Haimbaugh, Patricia H. Janak, Jocelyn M. Richard

## Abstract

A critical area of inquiry in the neurobiology of alcohol abuse is the neural mechanisms by which cues gain the ability to elicit alcohol use. We previously showed that cue-evoked activity in rat ventral pallidum (VP) robustly encodes the value of cues trained under both Pavlovian and instrumental contingencies, despite a stronger relationship between cue-evoked responses and behavioral latency after instrumental training. Here, we assessed VP neural representations of cue value in rats trained with a Pavlovian conditioned stimulus (CS+) that predicted alcohol delivery, and in rats trained with an instrumental discriminative stimulus (DS) that predicted alcohol availability if the rat entered the reward port during the cue. We also examined the impact of alcohol exposure itself on the integrity of this type of signaling in rats trained with sucrose. Decoding of cue value based on VP firing was blunted for an alcohol CS+ versus an alcohol DS, as well as in comparison to a sucrose DS or CS+. Further, homecage alcohol exposure had opposing effects on VP encoding of cue value for a sucrose DS versus a sucrose CS+, enhancing decoding accuracy for the DS and reducing decoding accuracy for the CS+. These findings suggest that problem alcohol seeking may result from biased engagement of specific reward-related processes via changes in VP signaling.

## Introduction

Addiction to alcohol and other drugs has been characterized as a shift away from decision-making based on representations of outcome value (goal-directed or ‘model-based’ reward seeking) toward stimulus evoked reward-seeking based on either the incentive motivational properties of cues (Robinson and Berridge, 2000), the development of habits (Everitt and Robbins, 2005; Vandaele and Janak, 2018) or ‘model-free’ reward-seeking (Lucantonio et al., 2014; Sebold et al., 2014; Reiter et al., 2016). Different classes of drugs may selectively disrupt different aspects of decision-making (Sweis et al., 2018). Further, associative and non-associative consumption may exert dissociable effects on decision-making. For instance, non-associative alcohol exposure has been shown to enhance the incentive motivational properties of cues (Spoelder et al., 2015; Kruse et al., 2017) and increase habit-like responding (Corbit et al., 2012; Corbit and Janak, 2016a; Houck and Grahame, 2018). Thus, it is critical that we determine not just what types of associative processes are engaged by alcohol as a reinforcer, but also the effects non-associative alcohol exposure has on brain mechanisms of learning and decision-making.

The ventral pallidum (VP) has been suggested to act as part of a common pathway for relapse to drug seeking, but also contributes to basic affective processes critical for normal well-being (Smith et al., 2009; Creed et al., 2016; Heinsbroek et al., 2017). The VP is implicated in promoting reward-seeking actions elicited by drug and alcohol-associated cues (Farrell et al., 2018), including renewal of operant responding for alcoholic beer by a beer-associated context (Perry and McNally, 2013; Prasad and McNally, 2016) and reinstatement of responding for cocaine by contingent presentations of a cocaine cue (Mahler et al., 2014). Importantly, VP activation correlates with self-reported alcohol craving in humans (Kühn and Gallinat, 2011). Activity in VP neurons in response to reward-related cues has been found to encode both incentive motivation (Smith et al., 2011; Ahrens et al., 2016; Richard et al., 2016) and outcome value (Tindell et al., 2009). To determine how VP activity contributes to alcohol abuse and addiction, we must define the specific associative and decision-making processes that are mediated by populations of VP neurons, and how encoding in these neurons is altered in models of alcohol abuse.

Previously, we found that VP cue responses similarly encode the value of cues trained under Pavlovian versus instrumental contingencies (Richard et al., 2018). In the instrumental task, rats are trained with a cue (a discriminative stimulus-DS) that predicts reward availability *if* the rat makes an entry to the reward port during the cue. In the Pavlovian task, rats are trained with a cue (a conditional stimulus-CS) that predicts reward delivery to the port irrespective of the rat’s behavior. Across learning, rats in both tasks increase port entry behavior during DS or the CS cue, resulting in superficially similar behavior during the cue period in both tasks. Activity in VP neurons encodes the learned value of both the DS and the CS, despite the differing action requirements in the two tasks.

Here, we examined VP encoding of an alcohol DS trained through instrumental conditioning versus an alcohol CS trained via Pavlovian conditioning. We found selective disruption of VP encoding of cue identity for an alcohol CS, but not an alcohol DS, in comparison to previously reported encoding of sucrose DS and CS cue identities (Richard et al., 2018). Because our effects could be due to the associative or non-associative properties of alcohol, we next investigated the impact of non-associative alcohol exposure on VP encoding of DS and CS cues for sucrose reward. We found that alcohol exposure alone had opposing effects on encoding of these sucrose cues, enhancing encoding of the DS and disrupting encoding of the CS. Our results suggest associative and non-associative alcohol exposure can interact to potentiate and disrupt different forms of cue-elicited behaviors.

## Materials and Methods

### Subjects

Male and female Long Evans rats (n=18; Envigo) weighing 250-275 grams at arrival were individually housed in a temperature-and humidity-controlled colony room on a 12 h light/dark cycle. All experimental procedures were approved by the Institutional Animal Care and Use Committee at Johns Hopkins University and were carried out in accordance with the guidelines on animal care and use of the National Institutes of Health of the United States.

### Alcohol Pre-Exposure

Rats were pre-exposed to ethanol in the homecage prior to training as described previously (Simms et al., 2008; Remedios et al., 2014; Millan et al., 2017). This was conducted to either 1) acclimatize them to the taste and pharmacological effects of ethanol and allow stabilization of drinking behavior prior to conditioning with alcohol at the reward, or 2) to assess the impact of ethanol exposure itself, in rats that were subsequently conditioned with sucrose reward. After one week of continuous access to 10% ethanol in the home cage, rats received chronic intermittent access to 20% ethanol in the homecage, for 24 hours at a time on Monday, Wednesday, and Friday, for a period of 7 weeks. Rats with ethanol intake <2 g/kg/day during the last two weeks of pre-exposure were excluded.

### Pavlovian Conditioning

Rats (n=10) underwent Pavlovian conditioning with either 15% ethanol (n=6) or 10% sucrose (n=4) as the reward. Rats were randomly assigned one of the following auditory cues as their conditioned stimulus (CS+) for training and testing: 1) white noise 2) a 2900 Hz tone, or 3) a siren tone (ramped from 4 to 8 kHz with a 400ms period). Rats received an alternate auditory cue as their CS−. During ethanol conditioning sessions, the CS+ and CS−, each lasting 10s, were presented on a pseudorandom variable interval schedule with a mean inter-trial interval (ITI) of 80s. At 9 s after the CS+ onset, 0.065 mL of 15% ethanol was delivered into the reward delivery port over a period of 1s. Rats underwent conditioning every other day until they met final response criteria (port entries on at least 70% of CS+ presentations and less than 30% of CS− presentations) prior to being implanted with electrode arrays. During sucrose conditioning sessions, the CS+ and CS−, each lasting 10s, were presented on a pseudorandom variable interval schedule with a mean inter-trial interval (ITI) of 35s. At 8s after the CS+ onset, 0.13 mL of 10% sucrose was delivered into the reward delivery port over a period of 2s (Richard et al., 2018). Rats underwent daily conditioning (sucrose) until they met final response criteria (port entries on at least 70% of CS+ presentations and less than 30% of CS− presentations) prior to being implanted with electrode arrays.

### DS task training

Rats (n=8) were trained to perform a modified DS task, similar to described previously (Ghazizadeh et al., 2012; Richard et al., 2016, 2018), with either 15% ethanol (n=5) or 10% sucrose (n=3) as the reward. Rats in the modified DS task were randomly assigned one of the following auditory cues as their DS for training and testing: 1) white noise 2) a 2900 Hz tone, or 3) a siren tone (ramped from 4 to 8 kHz with a 400ms period). Rats received an alternate auditory cue as their NS (neutral stimulus). Entries into the reward delivery port during the DS presentation resulted in delivery of 15% ethanol (0.065 ml) or 10% liquid sucrose (0.13 ml) delivery and termination of the DS cue. Port entries during the NS presentation or during the inter-trial interval had no programmed consequences. Rats underwent the following sequential training stages 1) DS only, up to 60s in duration, 2) DS only, up to 30 s, 3) DS only, up to 20 s, 4) DS only up to 10 s, and 5) final stage, DS and NS. Once rats met preliminary DS response criteria at each stage (port entries on at least 60% of DS presentations) they were advanced to the next stage on the following day. During the final stage of the DS task, the DS and NS were presented on a pseudorandom variable interval schedule with a mean ITI of 80s in ethanol rats and 35s in sucrose rats, and each cue lasted up to 10s. Rats underwent training every other day (alcohol) or daily (sucrose) until they met final criteria (port entries on at least 70% of DS presentations and less than 30% of NS presentations) prior to being implanted with electrode arrays. Rats failing to meet performance criteria within 25 sessions were excluded from further study, resulting in the exclusion of 1 rat trained with ethanol.

### Surgeries

During surgery, rats were anesthetized with isoflurane (5%) and placed in a stereotaxic apparatus, after which surgical anesthesia was maintained with isoflurane (0.5-2.0%). Rats received pre-operative injections of carprofen (5 mg/kg) and topical lidocaine for analgesia, and cefazolin (75 mg/kg) to prevent infection. Guide cannulae, electrodes and microdrives were secured to the skull with bone screws and dental acrylic. All rats were given at least 7 days to recover prior to any tethering. After they had reached criteria in either task, rats (Pavlovian conditioning with alcohol, n=6; DS task with alcohol, n=4; alcohol-exposed Pavlovian conditioning with sucrose, n=4; alcohol-exposed DS task with sucrose, n=3) received unilateral arrays of 16 electrodes each (0.004” tungsten wires arranged in a bundle) attached to microdrive devices that allowed the entire array to be lowered by 80 or 160 µm increments. Electrode arrays were targeted to VP at +0.0 mm AP, +2.4 mm ML, and −8.0 mm DV.

### Electrophysiologal Recordings

Electrophysiological recording was conducted as described previously (Nicola et al., 2004; Ambroggi et al., 2008; Ghazizadeh et al., 2010, 2012; Richard et al., 2016). Rats were connected to the recording apparatus (Plexon Inc, TX), consisting of a head stage with operational amplifiers, cable and a commutator to allow free movement during recording. Rats were run for 1.5 hr sessions every other day (ethanol) or 45 m sessions every day (sucrose). The microdrive carrying the electrode arrays was lowered by 80 to 160 µm at the end of each session with satisfactory behavior (>60% of DS or CS+ presentations with a response) in order to obtain a new set of neurons for each session that was included in the analysis. Sessions with unsatisfactory behavior or poor recordings were excluded from further analysis. Based on these criteria 2 rats implanted after Pavlovian conditioning with ethanol and one rat implanted after Pavlovian conditioning with alcohol and sucrose were excluded from further analysis.

### Analysis of Electrophysiological Recordings

#### Spike sorting

Isolation of individual units was performed off-line with Offline Sorter (Plexon) using principal component analysis as described previously (Ambroggi et al., 2008; Ghazizadeh et al., 2012). Interspike interval distribution, cross-correlograms and autocorrelograms were used to ensure that single units were isolated. Only units with well-defined waveforms with characteristics that were constant over the entire recording session were included in the study. Sorted units were exported to NeuroExplorer 3.0 and Matlab for further analysis.

#### Response detection

Peri-stimulus time histograms (PSTHs), constructed around the behavioral events using a bin size of 20 ms, were used to visualize firing during task events. The firing rate of each neuron during each bin of the PSTH was smoothed using a Lowess function and transformed to a z-score as follows: (F_i_ – F_mean_)/F_sd_, where F_i_ is the firing rate of the *i*th bin of the PSTH, and F_mean_ and F_sd_ are the mean and the SD of the firing rate during the 10s baseline period. Color-coded maps and average traces were constructed based on these z-scores. PSTHs were used to detect the presence of excitations and inhibitions, as well as their onsets and offsets, in comparison to a 10s baseline period prior to cue onset. At least one bin outside of the 99% confidence interval of the baseline during the analysis window for each event was required to determine an excitation or inhibition for that event (Richard et al., 2016, 2018). Because inhibitions are difficult to detect with this method (Richard et al., 2016), we also ran t-tests on firing rates during previously determined response windows timed to cue onsets (0-300 ms), post-cue port entries (−500 to 500 ms) and reward delivery times (0 to 2 s) in comparison to firing during the 10s pre-cue baseline period. For analysis of cue responses, we focused on the 300 ms post-cue window, as it occurs prior to the majority (85-90%) of post-cue movement onsets (Richard et al., 2018). To compare firing in response to different task events (e.g. DS versus NS), t-tests were run on event related firing during these same windows. To assess differences in population encoding (i.e. the degree to which the proportions of neurons that responded to task events differed) across the two tasks, we used chi-squared tests.

#### Receiver operating characteristic (ROC) Analysis

To additionally assess the ability of VP neural firing to predict the presence or identity of cues and subsequent behavioral responses, we conducted receiver operating characteristic (ROC) analysis evaluating either a) the DS window versus the NS window or b) the CS+ window versus the CS− windows. For each analysis, we assessed the probability that firing during each window met criteria that ranged from zero to the maximum firing rate for that neuron, and plotted the true positive rate (likelihood that the firing in the window of interest was above criteria) versus the false positive rate (likelihood that the firing in the control window was above criteria) to create a ROC curve for each neuron. We then assessed the area under the ROC curve (auROC) for all VP neurons as well as the average ROC curve for the whole population. To assess differences in auROC distributions across the task, reward and exposure conditions we used linear mixed effects models with task and reward or exposure as fixed effects and subject included as a random effect.

#### Decoding analysis

For single unit decoding, a linear discriminant analysis (LDA) model (the “fitcdiscr” function in Matlab) was trained on each neuron’s spike activity for 300 ms bins from 900ms pre-cue to 3s post-cue on 80% of trials. This model was then used to classify the remaining 20% of trials as reward cue or control cue (DS versus NS or CS+ versus CS−). We performed this 5 times in a 5-fold cross-validation approach and averaged performance across all 5 repetitions to find each unit’s accuracy. We also conducted the analysis with the trial identities shuffled to determine the accuracy on shuffled data. We then repeated this analysis for every neuron in each region for each bin. We focused our statistical analysis on the 300ms post-cue, using linear mixed effects models to assess the impact of task, outcome and/or alcohol exposure on single unit accuracy, and to compare accuracy between true and shuffled data for each group. To look at how model classification accuracy increased with additional units, we pooled together separately recorded units. We found the 5-fold cross-validated accuracy for models trained on the activity of randomly selected pseudoensembles of 1, 5 10, 25, 50, and 100 units from each group during the first 300ms post-cue. For each level, we performed the analysis 20 times. To assess differences in ensemble decoding across the tasks we used 3-way ANOVAs with factors for task, reward or exposure condition, and ensemble size, followed by Tukey post-hoc tests.

### Histology

Animals were deeply anesthetized with pentobarbital and electrode sites were labeled by passing a DC current through each electrode. All rats were perfused intracardially with .9% saline following by 4% paraformaldehyde. Brains were removed, post-fixed in 4% paraformaldehyde for 4-24 hrs, cryoprotected in 25% sucrose for >48hrs, and sectioned at 50um on a microtome. We stained the tissue with cresyl violet and verified the location of recording sites using light microscopy. The dorsoventral location of recording sites was determined by subtracting the distance driven between recording sessions from the final recording location. Units recorded during sessions when the recording sites were determined to be localized outside of the VP were excluded from analysis.

## Results

To assess the degree to which VP neurons encode alcohol cues trained on different types of associations, we trained rats in either a cued instrumental task (the ‘DS task’) or in Pavlovian conditioning, both with 15% ethanol as the reward. In the DS task, entry into the reward port during the DS (an auditory cue lasting up to 10 s) resulted in delivery of 15% ethanol and termination of the DS cue, whereas port entries during a control auditory cue (the non-rewarded stimulus [NS]) or during the intertrial interval had no programmed consequences. For rats trained in Pavlovian conditioning, one auditory cue (the CS+) predicted delivery of ethanol to the reward port at the end of the cue, irrespective of the animal’s behavior during the cue. Presentations of a control auditory cue (the CS−) did not predict ethanol delivery. Rats were trained in one of these tasks for up to 25 sessions until they made port entries during at least 70% of alcohol cue presentations (DS or CS+) and a maximum of 30% of control cue trials (NS or CS−). One rat trained in the DS task was excluded after failing to reach criteria within 25 sessions.

By the end of training, rats were more likely to enter the reward port during the alcohol cue (DS or CS+) then during the control cue (Figure 1A-B; main effect of cue F(1,8)=639.3, p=6.411e-9) regardless of task (interaction of task and cue F(1,8)=0.557, p=0.478), though port entry probability was higher in general during the DS task cues than during the Pavlovian cues (Figure 1F; main effect of task F(1,8)=15.10, p=0.0046). The impact of cue identity (reward versus control) on port entry latency differed based on task (interaction of task x cue F(1,8)=11.60, p=0.0093), due to a significant difference between port entry latency following the DS versus NS, but not for the CS+ versus CS− (Figure 1C-D, G). At the end of training, port entry latencies were shorter in the DS task versus the Pavlovian (Figure 1G; F(1,8)=71.58, p=2.9e-5). Rats trained in the DS task required more days to reach criteria (t(8)=2.855, p=0.02; Figure 1E).

**Figure 1.**
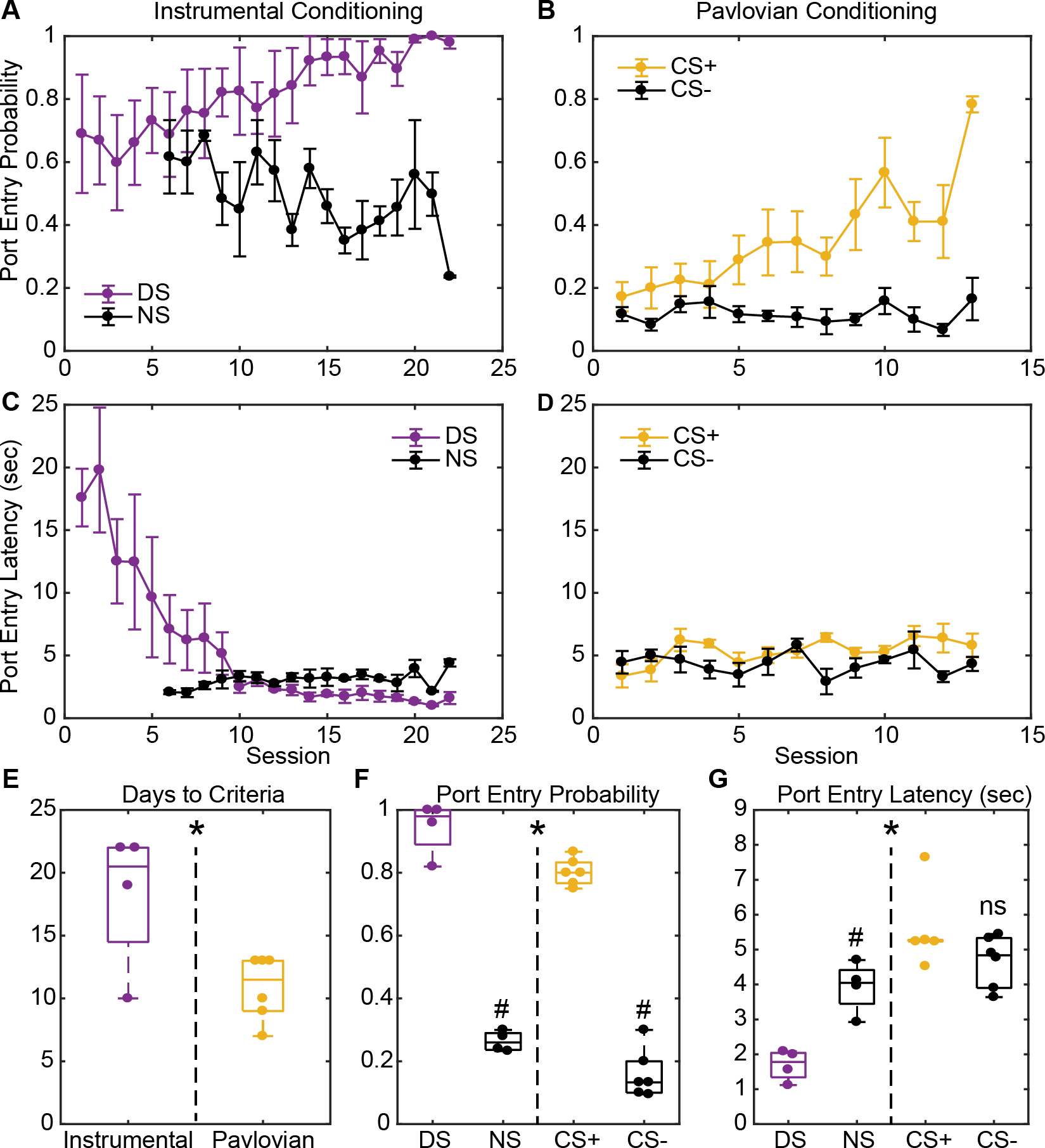
Summary of training behavior in the instrumental and Pavlovian tasks. Mean (+/− SEM) probability of port entry during the instrumental DS (A, purple) or the Pavlovian CS+ (B, yellow), and during the control cues (NS or CS−, black) across training until rats met response criteria. Mean (+/− SEM) latency to enter the port during the instrumental DS (C, purple) or the Pavlovian CS+ (D, yellow), and during the control cues (NS or CS−, black) across training until rats met response criteria for electrode implantation. Individual data points and mean +/− SEM for E) days to reach response criteria, F) final port entry probability, and G) final port entry latency (sec) prior to electrode array implantation. * = instrumental versus Pavlovian task difference, p < 0.05, # = alcohol versus control cue difference, p < 0.05, ns = non-significant difference for CS+ versus CS−

Once rats met training criteria, they were implanted with drivable electrode arrays aimed at the VP. Single unit recordings made during task performance were only included in task analysis if rats made port entries on at least 50% of alcohol cue presentations and less than 30% of control cue presentations, and if the response ratio ([probability of response to the alcohol cue]/[probability of response to the control cue + probability of response to the alcohol cue]) was at least 0.7. This resulted in exclusion of 2 rats from the Pavlovian conditioning group, and the inclusion of 126 units recorded during the DS task (4 rats, 14 sessions; Figure 2A) and 108 units recorded during Pavlovian conditioning (4 rats, 9 sessions; Figure 2B). Overall, many VP neurons were responsive to the alcohol cues and other tasks events (Figure 2C and D). Because we were primarily interested in cue encoding, we focused the bulk of our analysis on cue responses during the first 300 msec after cue onset.

**Figure 2.**
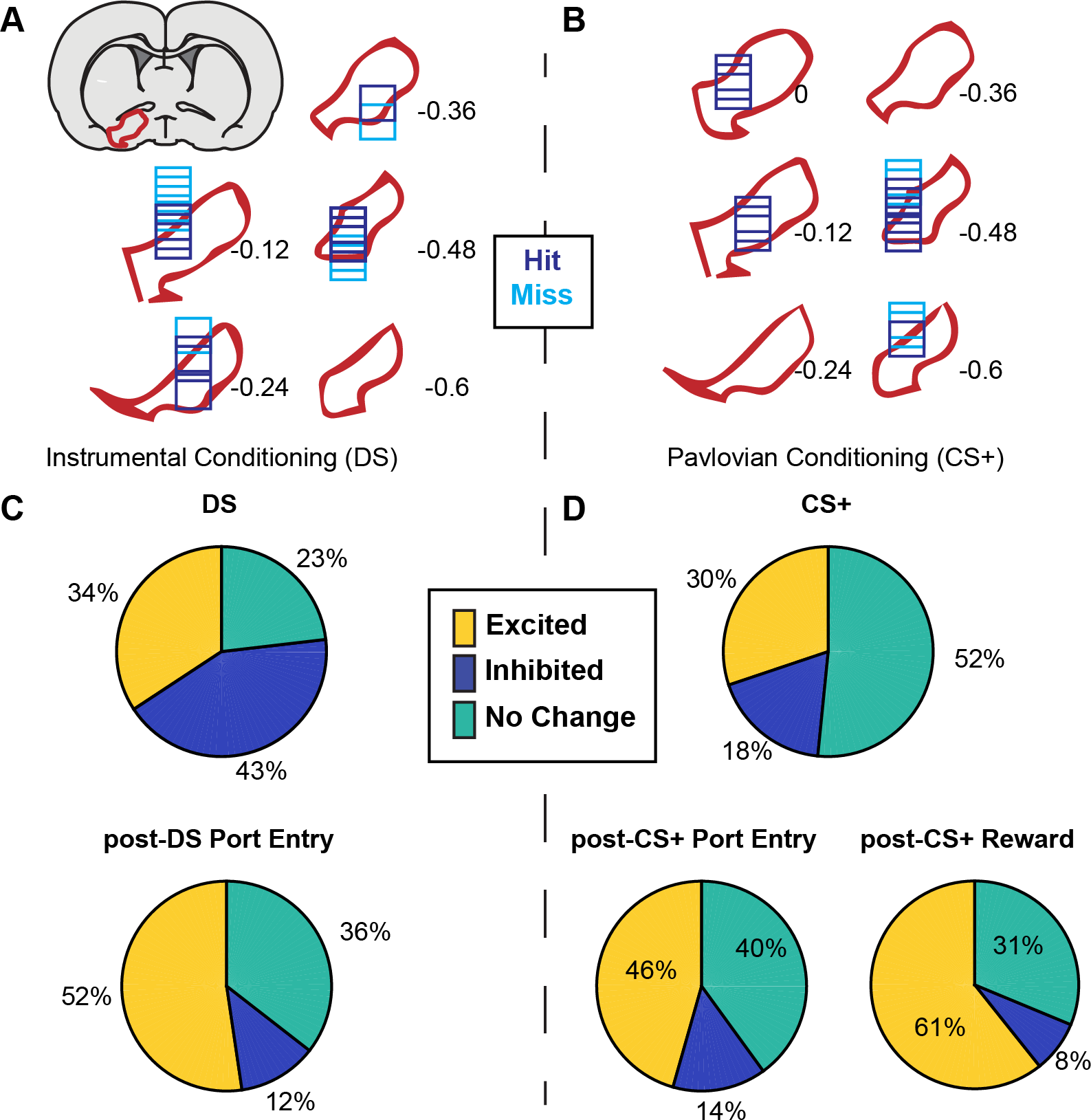
VP neurons respond to cues and reward seeking in both tasks. A) Histological reconstruction of electrode array placements in the instrumental task, shown on coronal slices, marked relative to bregma (mm), with the boundaries of VP demarcated in red. Purple squares mark array locations contained within VP, and blue squares mark array locations that were not contained within VP. B) Electrode array placements in the Pavlovian task. C) Pie charts showing the proportion of neurons (labels indicated percentage of units) for which we detected an increase (yellow) or decrease (dark blue) in firing following the DS and post-DS port entry D) Proportion of responsive neurons for CS+, post-CS+ port entry or post-CS+ reward delivery response windows.

### Ventral pallidal value encoding is more robust for a discriminative stimulus than for a Pavlovian cue for alcohol

Though the proportion of neurons responsive to cues was smaller than we previously reported for similar sucrose cues (Richard et al., 2016, 2018), similar proportions of neurons were excited by the two types of alcohol cues. Of units recorded in VP ~34% (38/108) were excited by the DS, and ~30% (38/126) were excited by the CS+ (Figure 2C-D; χ^2^ = 0.59; p=0.442). In contrast, a significantly smaller proportion of the population was inhibited by the alcohol CS+ (23/126, 18.2%) than by the alcohol DS (46/108, 42.6%; χ^2^ =16.58, p=4.7e-5). While similar proportions of neurons were excited by both cues type, the degree to which these responses were selective for the alcohol cue versus the control cue differed across the two tasks (Figure 3). Most DS-excited neurons (34/38) were significantly more excited by the DS than the NS (Figure 3A-B), and the DS-excited populations as a whole was significantly more excited by the DS than the NS (t(37)=8.897, p=1.149e-10; Figure 3E). In contrast, only 7 VP neurons were significantly more excited by the CS+ than the CS− (5% of recorded neurons, 18% of CS+ excited neurons; Figure 3C-D), though as a population CS+-excited neurons were significantly more excited by the CS+ than the CS− (t(37)=4.6677, p=3.92e-05; Figure 3G). For inhibitions, only 3 units were significantly more inhibited by the DS than the NS, though as a population DS-inhibited neurons were more inhibited by the DS than the NS (t(45)=−4.2702, =9.96e-05; Figure 3F). While only 6 units were more inhibited by the CS+ than the CS− (Figure 3D), CS+ inhibited neurons were more inhibited by the CS+ than the CS− on average (t(20)=−5.3934, p=2.8e-05; Figure 3H).

**Figure 3.**
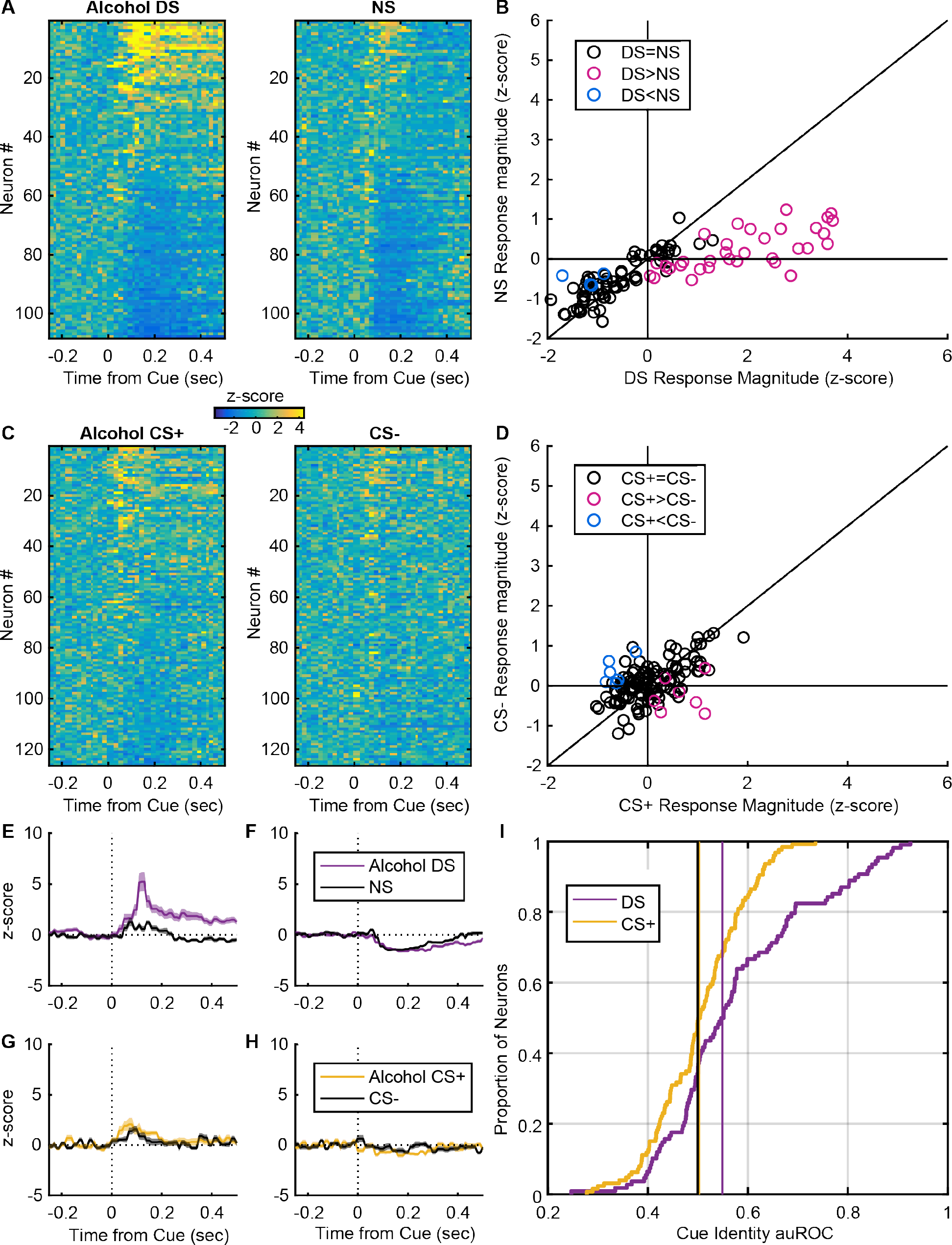
VP neurons encode the value of discriminative stimuli for alcohol availability but fail to encode the value of Pavlovian alcohol cues. A) Heatmaps of responses to the DS and NS from all neurons, sorted by the magnitude and direction of their response to the DS. Each line is the color coded, normalized (z-score) PSTH of an individual neuron. B) Scatterplot of normalized (mean z-score) responses of individual neurons to the DS versus the NS, showing neurons that fire significantly more following the DS (pink), the NS (blue) or neither cue (black). C) Heatmaps of all neurons’ responses to the CS+ and CS−, sorted by the magnitude and direction of their response to the CS+. Each line is the normalized (z-score), color coded PSTH. D) Scatterplot of normalized (mean z-score) responses to the CS+ versus the CS−, showing neurons that fire significantly more following the CS+ (pink), the CS− (blue) or neither cue (black). Average (mean +/− SEM) normalized response to the DS (purple) and NS (black) in DS excited (E) or DS inhibited (F) neurons. Average (mean +/− SEM) normalized response to the CS+ (yellow) and CS− (black) in CS+ excited (G) or CS+ inhibited (CS−) neurons. I) Cumulative distribution of auROCs for the assessment of encoding of cue identity for the DS versus the NS (purple) and the CS+ versus the CS− (yellow).

To quantify the degree to which increases in VP firing predicted the cue identity in the two tasks, we conducted receiver operating characteristic (ROC) analyses to assess the detection of the DS or the CS+ versus their respective control cues. Previously, we we found similar predictive encoding of cue identity for a sucrose DS and CS+ using this analysis (Richard et al., 2018). For each unit, we calculated the area on the ROC curve (auROC); auROCs greater than 0.5 indicate units that had greater firing rates on alcohol cue than control cue trials. We found significantly greater auROCs for the DS than the CS+ (Figure 3I; F(1,230)=77.751, p=2.949e-16), indicating that higher firing rates are more predictive of cue identity in the DS task than the CS+ and that excitatory encoding of cue value after Pavlovian conditioning is weaker.

### VP units and ensembles decode Pavlovian alcohol cues less accurately than both discriminative stimuli for alcohol and sucrose cues of either type

The previous analysis indicates weaker excitatory encoding of cue identity after Pavlovian conditioning with alcohol, versus training with a discriminative stimulus. One shortcoming of the ROC analysis was that it only assessed predictions of cue identity based on increases in firing, our *a priori* model of how VP neurons encode cue value. To examine possible differences in cue identity in spike activity of VP neurons, taking into account both increases and decreases in activity, we used we used linear discriminant analysis (LDA), an approach we used previously to decode reward identity based on VP firing (Ottenheimer et al., 2018). We trained LDA models based on the spike activity of individual neurons, and then, using 5-fold cross-validation, determined the accuracy of predicted cue identity (reward cue versus control cue). We focused our statistical comparisons across tasks on the first 300 msec after cue onsets. The decoding accuracy for cue identity in the alcohol DS task was significantly greater than decoding accuracy for the alcohol CS+ in comparison to shuffled control data (cue type [DS versus CS+] X shuffled: F(1,452)=5.49, p = 0.0195; Figure 4A-B). To compare decoding across sucrose and alcohol cues we reanalyzed previous recordings from rats performing these tasks with sucrose (Richard et al., 2018) using the LDA approach. When we assessed the accuracy of single unit decoding of cue identity across these tasks, we found that decoding for the alcohol CS+ was significantly less accurate than either the alcohol DS or the two sucrose cues (Figure 4A-E; interaction of cue type [DS versus CS+] and reward [alcohol versus sucrose], F(1,798)=5.07, p=0.0256; effect of cue type in alcohol trained rats, F(1,225)=28.926, p=1.88e-7).

**Figure 4.**
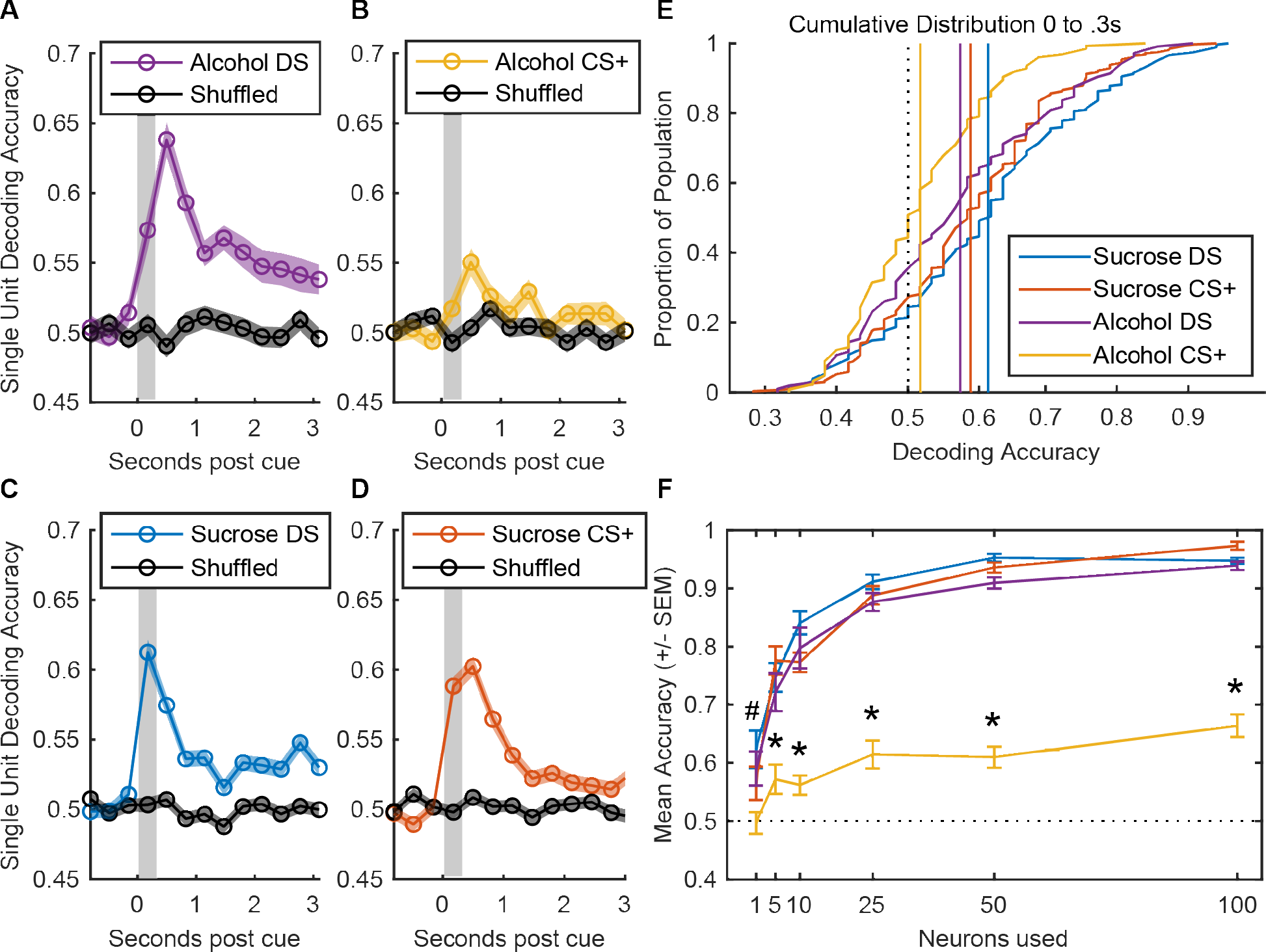
Decoding of alcohol-predicting Pavlovian cues based on VP activity is compromised relative to instrumental cues and sucrose-rewarded tasks. Average (mean +/− SEM) accuracy decoding cue identity with linear discriminant models trained on VP singe-unit activity at the time of the predictive and non-predictive cues for the instrumental (A,C) and Pavlovian (B,D) tasks with sucrose (A,B) and alcohol (C,D) as the rewards. Accuracy is plotted for 300ms bins relative to cue onset for both true data and data with cue identity shuffled. E) Cumulative distribution of unit model accuracies for each task and reward combination for the first 300ms following cue onset (gray shading in A-D). F) Average (mean +/− SEM) accuracy decoding cue identity with models trained on pseudoensembles of VP units from each task reward combination, ranging in size from 1 to 100 units. # = alcohol CS+ < alcohol DS and sucrose CS, * = alcohol CS+ < all other cues, p < 0.05 post-hoc Tukey tests.

The results so far suggest weaker encoding of alcohol CS+ cues. One possibility is that encoding of alcohol CS+ identity or value is more distributed across neurons. To assess this we pooled neurons together into pseudoensembles to compare how much information is contained within larger groups of VP neurons across each task. We then trained LDA models on the spiking activity of 5, 10, 25, 50 and 100 units randomly selected from the total recorded population. Increasing the number of neurons increased model accuracy (Figure 4F; F(5,456)=159.81, p=7.47e-98), with more improvement in accuracy occurring for the alcohol DS and sucrose cues (Figure 4F; interaction of ensemble size X cue type X reward, F(5,456)=7.9635, p=3.286-07). Overall decoding accuracy remained blunted for the alcohol CS+, even at larger ensemble sizes (interaction of cue type X reward, F(1,456)=139.18 p=3.228e-28). While decoding of the alcohol DS or sucrose cues versus their respective control cues reached over 90% accuracy at the largest ensemble size, mean accuracy for the alcohol CS+ remained less than 65%.

### Alcohol exposure modifies pallidal encoding of conditioned and instrumental stimuli in a diverging fashion

Our observation that representations of Pavlovian cue identity by VP firing was compromised in alcohol sessions relative to sucrose sessions, while the encoding of the DS versus NS was largely intact, led us to question how alcohol affects these two tasks differently. Importantly, the alcohol experimental protocol involves two manipulations—a non-associative exposure period of intermittent access in the home cage, and a subsequent associative use of alcohol as the reward outcome during the task—so we were interested in testing which aspect of the rats’ experience with alcohol consumption was responsible for the divergent cue encoding. To this end, we exposed a new group of rats to homecage alcohol but then trained them in Pavlovian conditioning or in the DS task with sucrose as the reward. This approach permitted exploration of whether alcohol exposure alone changes encoding of CS+ and DS.

Unexpectedly, alcohol exposure had a profound effect on VP responses to cues. For the DS, 88% of neurons were cue-excited, an even greater proportion than the 60% we found previously for rats without alcohol exposure performing the DS task for sucrose reward; for the CS+, 44% of neurons were cue-excited, comparable to the 49% in non-exposed animals (Richard et al., 2018). In alcohol-exposed rats trained on the instrumental task, a large proportion of VP neurons had greater firing for the DS than for the NS (49%, 82/166; Figure 5A-B). Fewer VP neurons from alcohol-exposed rats trained on the CS+ task had greater firing for the CS+ than the CS− (28%, 47/170; Fig. 5C-D). While cue responsive neurons (aside from the 6 DS-inhibited neurons) responded on the whole significantly differently to the sucrose versus control cue in both tasks (DS-excited: t(115)=11.27, p=2.663e-20; DS-inhibited: t(5)=−1.529, p=0.1868; CS+-excited: t(44)=11.95, p=2.08e-15; CS+-inhibited: t(25)=−6.637, p=5.915e-7; Figure 5E-H), cue identity prediction based on increases in firing following cue onset was significantly greater on average for the sucrose DS versus the sucrose CS+ (F(1,334)=8.064, p=0.00479; Fig. 5I). This finding differed from our previous comparison of recordings during sucrose DS and CS+ cues in rats not exposed to alcohol (Richard et al., 2018), which showed no difference in auROC distributions for cue discriminability based on increases in neural firing.

**Figure 5.**
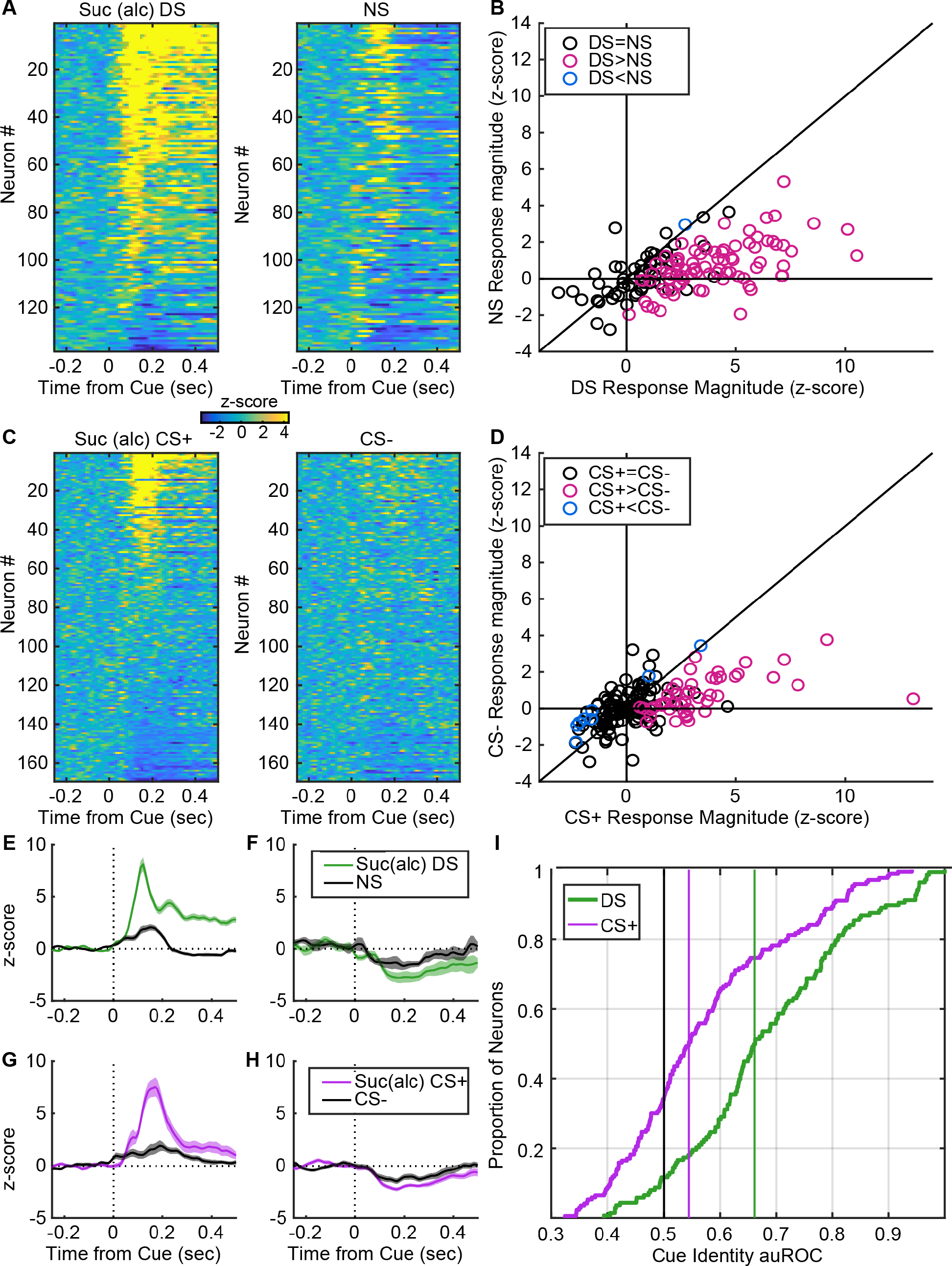
Following home cage alcohol exposure, VP neural responses are more predictive of the learned value of discriminative stimuli than Pavlovian cues, even with sucrose reward. A) Heatmaps of responses to the DS and NS from all neurons, sorted by the magnitude and direction of their response to the DS. Each line is the normalized (z-score), color coded PSTH of an individual neuron. B) Scatterplot of normalized (mean z-score) responses to the DS versus the NS, showing neurons that fire significantly more following the DS (pink), the NS (blue) or neither cue (black). C) Heatmaps of responses to the CS+ and CS− from all neurons, sorted by the magnitude and direction of their response to the CS+. Each line is the normalized (z-score), color coded PSTH of an individual neuron. D) Scatterplot of normalized (mean z-score) responses to the CS+ versus the CS−, showing neurons that fire significantly more following the CS+ (pink), the CS− (blue) or neither cue (black). Average (mean +/− SEM) normalized response to the DS (green) and NS (black) in DS excited (E) or DS inhibited (F) neurons. Average (mean +/− SEM) normalized response to the CS+ (pink) and CS− (black) in CS+ excited (G) or CS+ inhibited (CS−) neurons. I) Cumulative distribution of auROCs for the assessment of encoding of cue identity for the DS versus the NS (green) and the CS+ versus the CS− (pink).

As before, this initial approach specifically tested whether neurons had greater firing for the rewarded cue over the non-rewarded cue. To evaluate the robustness of cue encoding in VP in each task more neutrally, and to better compare these results to those from non-exposed rats, we used single-unit and pseudoensemble decoding methods (Figure 6). Consistent with our initial results, the cue discrimination accuracy of single-unit models trained on neurons from alcohol-exposed rats in the DS task improved over shuffled data more than those trained on neurons from alcohol-exposed CS+ rats (cue type X shuffled: F(1,636)=21.417, p=4.460e-6; Fig. 6A). Notably, decoding accuracy was shifted higher for alcohol-exposed DS rats relative to non-exposed DS rats, but not for CS+ rats (cue type X exposure: F(1,1786)=7.161, p=0.00752; Fig. 6B). When looking at the distribution of information across the population via pseudoensembles, we once again found divergent effects of alcohol exposure on discriminable cue encoding, seen in a significant interaction between the effects of task and alcohol exposure (F(1,456)=34.0, p=1.04e-8) that did not vary across ensemble size (interaction of ensemble size X cue type X alcohol exposure: F(5,456)=0.759, p=0.580; Fig. 6C-D). Moreover, post-hoc Tukey tests revealed that, on the whole, models trained on ensembles from alcohol-exposed DS rats were significantly more accurate than those from non-exposed DS rats (p=0.0000533); in contrast, models trained on alcohol-exposed CS+ rats were less accurate than those from non-exposed CS+ rats (p=0.000809). The data indicate that alcohol exposure has opposing effects on cue value encoding depending on the associative structure of the task.

**Figure 6.**
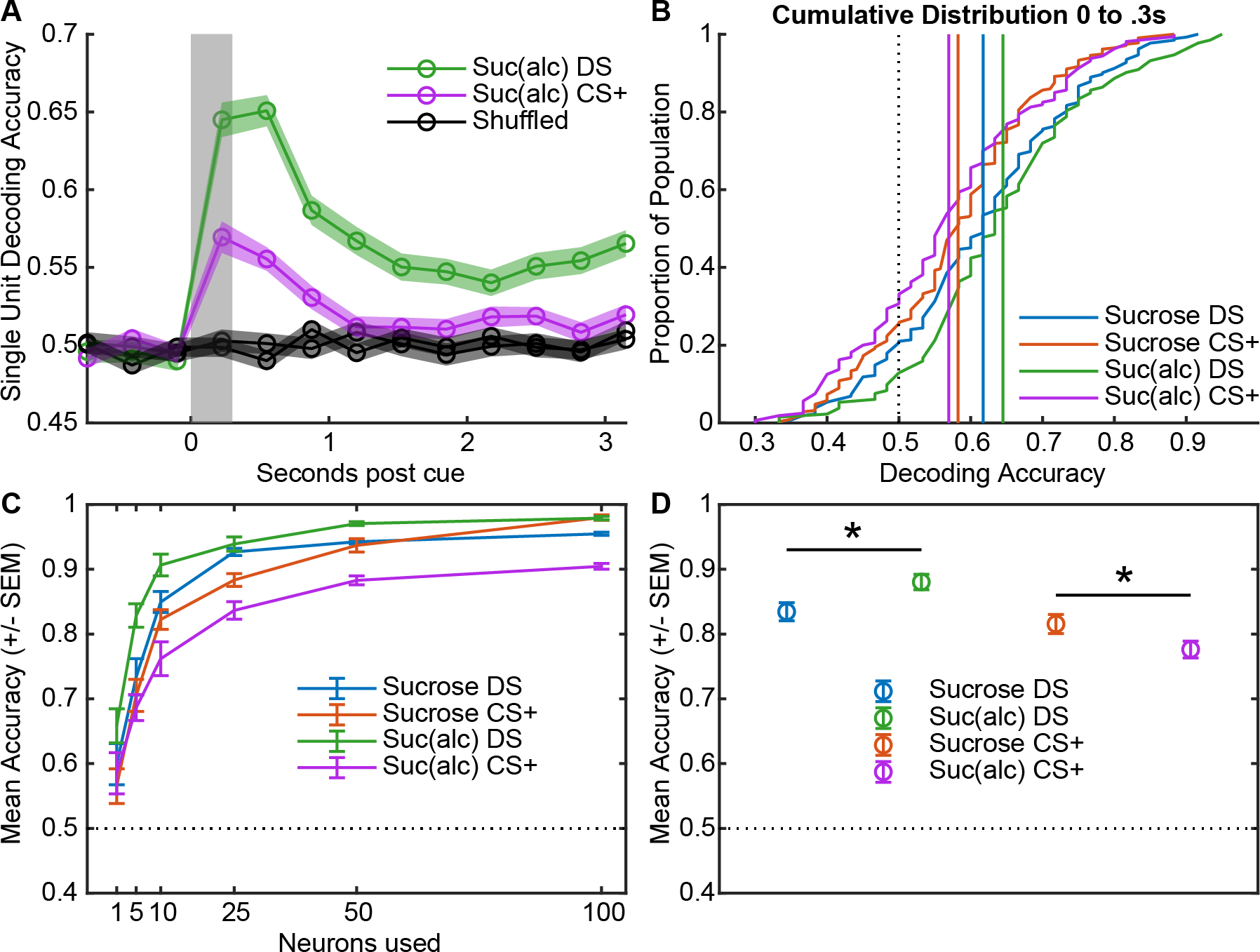
Alcohol exposure blunts decoding accuracy of Pavlovian cues while enhancing decoding of instrumental cues. A) Average (mean +/− SEM) accuracy decoding cue identity with linear discriminant models trained on VP singe-unit activity at the time of the predictive and non-predictive cues for the instrumental (green) and Pavlovian (purple) tasks with sucrose as the reward in rats with previous homecage alcohol exposure. Accuracy is plotted for 300ms bins relative to cue onset for both true data and data with cue identity shuffled (black). D) Cumulative distribution of unit model accuracies for each task and exposure combination the first 300ms following cue onset (gray shading in A) for all rats trained with sucrose reward. C) Average (mean +/− SEM) accuracy decoding cue identity with models trained on pseudoensembles of VP units from each task-exposure combination for rats trained on sucrose, ranging in size from 1 to 100 units. D) Average (mean +/− SEM) decoding accuracy across all ensemble sizes in (C). Asterisk indicates p < 0.001 for Tukey tests comparing the accuracy of the same task with different exposure conditions, corrected for multiple comparisons.

## Discussion

Here we find that VP responses to DS and CS cues are impacted differently by associative and non-associative alcohol exposure. When rats were trained with a DS or CS predicting alcohol (associative exposure), decoding of cue value is blunted for an alcohol CS, in comparison to an alcohol DS, a sucrose CS, or a sucrose DS. That is, VP neurons do not distinguish reliably between an alcohol CS cue and a control cue, even when rats reliably distinguish between these cues in their behavior. In contrast, VP neurons reliably distinguish between the reward cue and control cue in the other three paradigms (alcohol DS, sucrose DS, sucrose CS). Weaker encoding of the alcohol CS+ is not accounted for by lower numbers of cue-excited neurons, as we found a similar proportion of neurons were excited by an alcohol DS. Additionally, VP encoding of alcohol CS value is not simply more distributed throughout the neural population. When we attempted to decode the identity of each reward cue when it was presented based on models trained on increasingly large ensembles of VP neurons, we found that increasing the number of neurons included in the models improves performance for all cues more so than the alcohol CS+. Exposure to alcohol does not have to occur in an associative context for it to impact VP cue encoding. Voluntary exposure to alcohol in the homecage (non-associative exposure) has opposing effects on future VP encoding of cue value for a sucrose DS versus a sucrose CS at all ensemble sizes. Alcohol exposure enhances decoding accuracy for the DS and reduces decoding accuracy for the CS. These findings provide evidence that representations of DS versus CS associations by VP neurons may be differentially engaged in models of alcohol abuse.

### Disruptions of cue learning or encoding by alcohol exposure

Despite reported effects of alcohol on decision-making (Richards et al., 1999; Dougherty et al., 2008; Schindler et al., 2014; Irimia et al., 2015) and the importance of cues in driving alcohol use (Ludwig, 1986; Bordnick et al., 2008; Tomie and Sharma, 2013), studies characterizing the effects of alcohol exposure on neural encoding of reward-related cues have been limited. Ethanol exposure has been shown to disrupt encoding of immediate versus delayed rewards by nucleus accumbens neurons (Gutman and Taha, 2016). Chronic intermittent exposure to ethanol or training with an alcoholic food reward both separately enhance dopamine release to a Pavlovian CS, though combining these manipulations has no effect in comparison to controls (Fiorenza et al., 2018). Adolescent alcohol exposure has also been shown to enhance dopamine release to Pavlovian cues (Schindler et al., 2016). We are not aware of any previous investigations of ethanol effects on VP cue encoding, though ethanol injections into the ventral tegmental area have been shown to increase extracellular dopamine levels in VP (Ding et al., 2011), and systemic ethanol decreases levels of extracellular GABA in VP (Kemppainen et al., 2010). Opposing effects of alcohol on GABA and dopamine neurotransmitter release may play a role in the effects reported here.

### Opposing effects of alcohol exposure on encoding of CS versus DS cues

The low accuracy of cue discrimination models trained on VP activity during alcohol CS+ and CS− cues may be due to disruptions in VP encoding of cue value. Alternatively, reduced decoding accuracy for an alcohol CS+ may be due to alcohol itself being an outcome with low or mixed value. If alcohol is truly low in value, then why are VP responses to DS cues for alcohol relatively robust? One possibility is that DS cues elicit reward-related representations in VP neurons that are less tied to outcome value and more related to action value. For instance, we have previously shown that encoding of response latency by VP neurons is more robust after instrumental conditioning with a DS in comparison to Pavlovian conditioning with a CS cue (Richard et al., 2018).

Addiction has been characterized as a shift away from goal-directed behavior based on representations of the outcome value, toward cue-driven reward-seeking that occurs independently of expected outcome value (Everitt et al., 2001; Ostlund and Balleine, 2008; Corbit and Janak, 2016a). This shift leads to perseverative or compulsive drug-seeking despite adverse consequences. Conditioning with or exposure to alcohol may have opposing effects on the types of reward-related representations elicited by DS and CS cues. Extended training with or exposure to alcohol makes both alcohol and sucrose seeking behavior less sensitive to outcome devaluation (Corbit et al., 2012). In addition, acute alcohol exposure makes low-responding animals that are normally goal-directed less sensitive to outcome devaluation (Houck and Grahame, 2018). Cue-driven, outcome-devaluation-insensitive responses may be driven by the incentive motivational properties of cues (Robinson and Berridge, 2000). The incentive properties of cues lend them the power to attract attention and approach and to maintain or invigorate behaviors in the absence of the reward itself (Wyvell and Berridge, 2000; Cardinal and Everitt, 2004; Flagel et al., 2008). Alcohol exposure has been shown to enhance the incentive motivational properties of cues, including their ability to elicit approach and to reinforce novel behavioral responses (Spoelder et al., 2015; Kruse et al., 2017). Extended training with alcohol or sucrose increases the ability of Pavlovian cues for those rewards to invigorate separately trained instrumental responses (Corbit and Janak, 2016b).

The opposing effects of alcohol exposure on VP encoding of DS versus CS cues reported here may be due to differential representations of outcome value by VP responses to CS and DS cues. Because CS cues predict reward delivery independently of any explicit action contingencies, representations of outcome value by VP neurons may predominate after Pavlovian conditioning. Consistent with this, VP responses to CS cues predicting salt have been shown to track the current value of salt in the absence of opportunities for new learning (Tindell et al., 2009). Port entry responses to CS cues (sometimes referred to as ‘goal-tracking’), which are our behavioral measure of CS learning here, have been shown to be more sensitive to outcome devaluation than approach to the CS itself (i.e. ‘sign-tracking’), and may be more sensitive than behavioral responses to a DS (Colwill and Rescorla, 1990; Morrison et al., 2015; Vandaele et al., 2017). In this context, it may not be surprising that VP responses to Pavlovian cues are less robust after associative or non-associative exposure to alcohol, in comparison to DS cues.

## Conclusions

Overall, we find that VP encoding of the value of Pavlovian and instrumental cues differs when those cues are trained with alcohol as a reward or if rats are trained with the cues after alcohol exposure. Problem alcohol seeking is driven by biased engagement of specific reward-related processes, presumably due to altered signaling in reward-related circuitry. These results indicate that VP is one of these altered brain sites. The opposing effects of alcohol on encoding of these types of cues by VP neurons highlights the importance of better understanding the types of information that are signaled by VP neural responses to these cues, and how they are altered in models of drug and alcohol abuse.

## Acknowledgements

This work was supported by National Institutes of Health grants F32 AA022290 (JMR), K99/R00 AA025384 (JMR), R01 AA014925 (PHJ) and R01 AA026306 (PHJ), by a NARSAD Young Investigator Award (JMR), and by the National Science Foundation Graduate Research Fellowship under Grant No. DGE-1746891 (DO).

## References

Ahrens AM, Meyer PJ, Ferguson LM, Robinson TE, Aldridge JW (2016) Neural Activity in the Ventral Pallidum Encodes Variation in the Incentive Value of a Reward Cue. J Neurosci 36:7957–7970.

Ambroggi F, Ishikawa A, Fields HL, Nicola SM (2008) Basolateral amygdala neurons facilitate reward-seeking behavior by exciting nucleus accumbens neurons. Neuron 59:648–661.

Bordnick PS, Traylor A, Copp HL, Graap KM, Carter B, Ferrer M, Walton AP (2008) Assessing reactivity to virtual reality alcohol based cues. Addict Behav 33:743–756.

Cardinal RN, Everitt BJ (2004) Neural and psychological mechanisms underlying appetitive learning: links to drug addiction. Curr Opin Neurobiol 14:156–162.

Colwill RM, Rescorla RA (1990) Effect of reinforcer devaluation on discriminative control of instrumental behavior. J Exp Psychol Anim Behav Process 16:40–47.

Corbit LH, Janak PH (2016a) Habitual Alcohol Seeking: Neural Bases and Possible Relations to Alcohol Use Disorders. Alcohol Clin Exp Res 40:1380–1389.

Corbit LH, Janak PH (2016b) Changes in the Influence of Alcohol-Paired Stimuli on Alcohol Seeking across Extended Training. Front Psychiatry 7:169.

Corbit LH, Nie H, Janak PH (2012) Habitual alcohol seeking: time course and the contribution of subregions of the dorsal striatum. Biol Psychiatry 72:389–395.

Creed M et al. (2016) Convergence of Reinforcing and Anhedonic Cocaine Effects in the Ventral Pallidum. Neuron 92:119–146.

Ding Z-M, Oster SM, Hall SR, Engleman EA, Hauser SR, McBride WJ, Rodd ZA (2011) The stimulating effects of ethanol on ventral tegmental area dopamine neurons projecting to the ventral pallidum and medial prefrontal cortex in female Wistar rats: regional difference and involvement of serotonin-3 receptors. Psychopharmacology (Berl) 216:245–255.

Dougherty DM, Marsh-Richard DM, Hatzis ES, Nouvion SO, Mathias CW (2008) A test of alcohol dose effects on multiple behavioral measures of impulsivity. Drug Alcohol Depend 96:111–120.

Everitt BJ, Dickinson A, Robbins TW (2001) The neuropsychological basis of addictive behaviour. Brain Res Brain Res Rev 36:129–138.

Everitt BJ, Robbins TW (2005) Neural systems of reinforcement for drug addiction: from actions to habits to compulsion. Nat Neurosci 8:1481–1489.

Farrell MR, Schoch H, Mahler S V. (2018) Modeling cocaine relapse in rodents: Behavioral considerations and circuit mechanisms. Prog Neuro-Psychopharmacology Biol Psychiatry.

Fiorenza AM, Shnitko TA, Sullivan KM, Vemuru SR, Gomez-A A, Esaki JY, Boettiger CA, Da Cunha C, Robinson DL (2018) Ethanol Exposure History and Alcoholic Reward Differentially Alter Dopamine Release in the Nucleus Accumbens to a Reward-Predictive Cue. Alcohol Clin Exp Res 42:1051–1061.

Flagel SB, Akil H, Robinson TE (2008) Individual differences in the attribution of incentive salience to reward-related cues: Implications for addiction. Neuropharmacology.

Ghazizadeh A, Ambroggi F, Odean N, Fields HL (2012) Prefrontal cortex mediates extinction of responding by two distinct neural mechanisms in accumbens shell. J Neurosci 32:726–737.

Ghazizadeh A, Fields HL, Ambroggi F (2010) Isolating event-related neuronal responses by deconvolution. J Neurophysiol 104:1790–1802.

Gutman AL, Taha SA (2016) Acute ethanol effects on neural encoding of reward size and delay in the nucleus accumbens. J Neurophysiol 116:1175–1188.

Heinsbroek JA, Neuhofer DN, Griffin WC, Siegel GS, Bobadilla A-C, Kupchik YM, Kalivas PW (2017) Loss of Plasticity in the D2-Accumbens Pallidal Pathway Promotes Cocaine Seeking. J Neurosci 37:757–767.

Houck CA, Grahame NJ (2018) Acute drug effects on habitual and non-habitual responding in crossed high alcohol preferring mice. Psychopharmacology (Berl).

Irimia C, Wiskerke J, Natividad LA, Polis IY, de Vries TJ, Pattij T, Parsons LH (2015) Increased impulsivity in rats as a result of repeated cycles of alcohol intoxication and abstinence. Addict Biol 20:263–274.

Kemppainen H, Raivio N, Nurmi H, Kiianmaa K (2010) GABA and glutamate overflow in the VTA and ventral pallidum of alcohol-preferring AA and alcohol-avoiding ANA rats after ethanol. Alcohol Alcohol 45:111–118.

Kruse LC, Schindler AG, Williams RG, Weber SJ, Clark JJ (2017) Maladaptive Decision Making in Adults with a History of Adolescent Alcohol use, in a Preclinical Model, Is Attributable to the Compromised Assignment of Incentive Value during Stimulus-Reward Learning. Front Behav Neurosci 11:134.

Kühn S, Gallinat J (2011) Common biology of craving across legal and illegal drugs - a quantitative meta-analysis of cue-reactivity brain response. Eur J Neurosci 33:1318–1326.

Lucantonio F, Caprioli D, Schoenbaum G (2014) Transition from ‘model-based’ to ‘model-free’ behavioral control in addiction: Involvement of the orbitofrontal cortex and dorsolateral striatum. Neuropharmacology 76:407–415.

Ludwig AM (1986) Pavlov’s “bells” and alcohol craving. Addict Behav 11:87–91.

Mahler S V, Vazey EM, Beckley JT, Keistler CR, McGlinchey EM, Kaufling J, Wilson SP, Deisseroth K, Woodward JJ, Aston-Jones G (2014) Designer receptors show role for ventral pallidum input to ventral tegmental area in cocaine seeking. Nat Neurosci 17:577–585.

Millan EZ, Kim HA, Janak PH (2017) Optogenetic activation of amygdala projections to nucleus accumbens can arrest conditioned and unconditioned alcohol consummatory behavior. Neuroscience.

Morrison SE, Bamkole MA, Nicola SM (2015) Sign Tracking, but Not Goal Tracking, is Resistant to Outcome Devaluation. Front Neurosci 9:468.

Nicola SM, Yun IA, Wakabayashi KT, Fields HL (2004) Cue-Evoked Firing of Nucleus Accumbens Neurons Encodes Motivational Significance During a Discriminative Stimulus Task. J Neurophysiol 91:1840–1865.

Ostlund SB, Balleine BW (2008) On habits and addiction: an associative analysis of compulsive drug seeking. Drug Discov Today Dis Model 5:235–245.

Ottenheimer D, Richard JM, Janak PH (2018) Ventral pallidum encodes relative reward value earlier and more robustly than nucleus accumbens. Nat Commun 9:4350.

Perry CJ, McNally GP (2013) A role for the ventral pallidum in context-induced and primed reinstatement of alcohol seeking. Eur J Neurosci 38:2762–2773.

Prasad AA, McNally GP (2016) Ventral Pallidum Output Pathways in Context-Induced Reinstatement of Alcohol Seeking. J Neurosci 36:11716–11726.

Reiter AMF, Deserno L, Wilbertz T, Heinze H-J, Schlagenhauf F (2016) Risk Factors for Addiction and Their Association with Model-Based Behavioral Control. Front Behav Neurosci 10:26.

Remedios J, Woods C, Tardif C, Janak PH, Chaudhri N (2014) Pavlovian-conditioned alcohol-seeking behavior in rats is invigorated by the interaction between discrete and contextual alcohol cues: implications for relapse. Brain Behav 4:278–289.

Richard JM, Ambroggi F, Janak PH, Fields HL (2016) Ventral Pallidum Neurons Encode Incentive Value and Promote Cue-Elicited Instrumental Actions. Neuron 90:1165–1173.

Richard JM, Stout N, Acs D, Janak PH (2018) Ventral pallidal encoding of reward-seeking behavior depends on the underlying associative structure. Elife 7:e33107.

Richards JB, Zhang L, Mitchell SH, de Wit H (1999) Delay or probability discounting in a model of impulsive behavior: effect of alcohol. J Exp Anal Behav 71:121–143.

Robinson TE, Berridge KC (2000) The psychology and neurobiology of addiction: an incentive-sensitization view. Addiction 95:91–117.

Schindler AG, Soden ME, Zweifel LS, Clark JJ (2016) Reversal of Alcohol-Induced Dysregulation in Dopamine Network Dynamics May Rescue Maladaptive Decision-making. J Neurosci 36:3698–3708.

Schindler AG, Tsutsui KT, Clark JJ (2014) Chronic Alcohol Intake During Adolescence, but not Adulthood, Promotes Persistent Deficits in Risk-Based Decision Making. Alcohol Clin Exp Res 38:1622–1629.

Sebold M, Deserno L, Nebe S, Schad DJ, Garbusow M, Hägele C, Keller J, Jünger E, Kathmann N, Smolka M, Rapp MA, Schlagenhauf F, Heinz A, Huys QJM, Heinz A, Huys QJM (2014) Model-Based and Model-Free Decisions in Alcohol Dependence. Neuropsychobiology 70:122–131.

Simms JA, Steensland P, Medina B, Abernathy KE, Chandler LJ, Wise R, Bartlett SE (2008) Intermittent access to 20% ethanol induces high ethanol consumption in Long-Evans and Wistar rats. Alcohol Clin Exp Res 32:1816–1823.

Smith KS, Berridge KC, Aldridge JW (2011) Disentangling pleasure from incentive salience and learning signals in brain reward circuitry. Proc Natl Acad Sci U S A 108:E255–64.

Smith KS, Tindell AJ, Aldridge JW, Berridge KC (2009) Ventral pallidum roles in reward and motivation. Behav Brain Res 196:155–167.

Spoelder M, Tsutsui KT, Lesscher HMB, Vanderschuren LJMJ, Clark JJ (2015) Adolescent Alcohol Exposure Amplifies the Incentive Value of Reward-Predictive Cues through Potentiation of Phasic Dopamine Signaling. Neuropsychopharmacology.

Sweis BM, Redish AD, Thomas MJ (2018) Prolonged abstinence from cocaine or morphine disrupts separable valuations during decision conflict. Nat Commun 9:2521.

Tindell AJ, Smith KS, Berridge KC, Aldridge JW (2009) Dynamic computation of incentive salience: “Wanting” what was never “liked”. J Neurosci 29:12220–12228.

Tomie A, Sharma N (2013) Pavlovian Sign-Tracking Model of Alcohol Abuse. Curr Drug Abuse Rev 6.

Vandaele Y, Janak PH (2018) Defining the place of habit in substance use disorders. Prog Neuro-Psychopharmacology Biol Psychiatry 87:22–32.

Vandaele Y, Pribut HJ, Janak PH (2017) Lever Insertion as a Salient Stimulus Promoting Insensitivity to Outcome Devaluation. Front Integr Neurosci 11:23.

Wyvell CL, Berridge KC (2000) Intra-accumbens amphetamine increases the conditioned incentive salience of sucrose reward: enhancement of reward “wanting” without enhanced “liking” or response reinforcement. J Neurosci 20:8122–8130.

